# Bacteriophage P22 SieA mediated superinfection exclusion

**DOI:** 10.1101/2023.08.15.553423

**Authors:** Justin C. Leavitt, Brianna M. Woodbury, Eddie B. Gilcrease, Charles M. Bridges, Carolyn M. Teschke, Sherwood R. Casjens

**Affiliations:** School of Biological Sciences, University of Utah, Salt Lake City, UT 84112 USA; Green Raccoon Scientific, Gunlock UT 84733 USA; Department of Molecular and Cell Biology, University of Connecticut, Storrs, CT 06269, USA; York Structural Biology Laboratory, Department of Chemistry, University of York, York YO10 5DD, UK; Division of Microbiology and Immunology, Pathology Department, University of Utah School of Medicine, Salt Lake City, UT 84112 USA; Department of Civil and Environmental Engineering, University of Utah, Salt Lake City, UT 84112 USA; Department of Chemistry, University of Connecticut, Storrs, CT 06269 USA

**Keywords:** P22, bacteriophage, superinfection exclusion, ejection proteins, SieA

## Abstract

Many temperate phages encode prophage-expressed functions that interfere with superinfection of the host bacterium by external phages. *Salmonella* phage P22 has four such systems that are expressed from the prophage in a lysogen that are encoded by the *c2* (repressor), *gtrABC*, *sieA*, and *sieB* genes. Here we report that the P22-encoded SieA protein is the only phage protein required for exclusion by the SieA system, and that it is an inner membrane protein that blocks DNA injection by P22 and its relatives, but has no effect on infection by other tailed phage types. The P22 virion injects its DNA through the host cell membranes and periplasm via a conduit assembled from three “ejection proteins” after their release from the virion. Phage P22 mutants were isolated that overcome the SieA block, and they have amino acid changes in the C-terminal regions of the gene *16* and *20* encoded ejection proteins. Three different single amino acid changes in these proteins are required to obtain nearly full resistance to SieA. Hybrid P22 phages that have phage HK620 ejection protein genes are also partially resistant to SieA. There are three sequence types of extant phage-encoded SieA proteins that are less than 30% identical to one another, yet comparison of two of these types found no differences in target specificity. Our data are consistent with a model in which the inner membrane protein SieA interferes with the assembly or function of the periplasmic gp20 and membrane-bound gp16 DNA delivery conduit.

**HIGHLIGHTS:**

- Phage P22 SieA protein blocks DNA injection by P22-like phages
- SieA is an inner membrane protein
- Hybrid P22 phages with phage HK620 ejection proteins partially escape SieA exclusion
- SieA escape mutants of P22 alter the gp16 and 20 proteins that form the DNA ejection tube.

## INTRODUCTION

Temperate tailed bacteriophages can use one of two strategies for replication: a lytic cycle, which is a productive infection that releases progeny virions through host cell lysis, or a lysogenic cycle, which establishes a prophage state where the integrated phage DNA replicates in synchrony with the host chromosome without major detriment to the host cell. Although prophages do not express the genes required for the lytic cycle, they do express a few genes whose products often modify their bacterial host. A significant fraction of such “lysogenic conversion” genes act to protect the host bacterium from infection by other phages, which is called superinfection exclusion.

Prophage-expressed superinfection exclusion systems have been studied in numerous phages, and they include systems that block infecting phages at various stages of the lytic cycle (reviewed by 1, 2, 3). Some prophages inhibit DNA delivery into host cells by expressing proteins that modify bacterial surface phage receptors and thus prevent the adsorption of superinfecting phages. These include enzymes that alter surface polysaccharides (4, 5) and proteins that interfere with the accessibility of outer membrane proteins (6, 7). Other superinfection exclusion systems block phage development after the initial adsorption event. For example, prophage-encoded proteins can block DNA injection by unknown mechanisms (*e.g.*, *Escherichia coli* phage HK97 gp15 (8), *Streptococcus thermophilus* phage TP-J34 Ltp protein (9) and *Lactococcus lactis* phage Tuc2009 Sie_2009_ protein (10)). Yet other exclusion systems interfere with incoming phages at later stages of the replication cycle; for instance, abortive infection systems cause replication failure in mid lytic growth. In such cases the infected cell dies but does not release progeny viruses, thus preventing their spread (reviewed by 11).

*Salmonella enterica* phage P22 prophages interfere with superinfecting phages in at least four different ways. These include (i) the prophage c2 repressor protein that prevents lytic gene expression by homo-immune phages (12–14); (ii) the three GtrABC proteins that alter the *S. enterica* Typhimurium O-antigen surface polysaccharide to block adsorption by phages that utilize it as a receptor (5, 15, 16); and two superinfection exclusion systems mediated by the less well understood (iii) *sieA* and (iv) *sieB* genes (17–19). The SieB exclusion system causes cellular macromolecular synthesis to cease midway through the lytic cycle during superinfection by P22-like phages but not by P22 itself (19–21). The SieA protein blocks superinfection by P22-like phages (including P22 itself) at a very early stage after adsorption (19, 22–24).

Previous studies have shown that SieA does not affect the adsorption of superinfecting P22 phage virions (18, 23, 24) or the production of progeny after P22 prophage induction (24). SieA exclusion is not specific for the type of DNA in the infecting virion since generalized transduction by phage P22 is also greatly lowered by a *sieA^+^* P22 prophage in the recipient cell (19, 22–24). In a *sieA^+^* lysogen superinfecting P22 DNA is not cleaved by cytoplasmic host restriction endonucleases, and superinfecting P22 phage genes are not expressed (20). These findings suggest that the *sieA* block occurs early in the infection cycle after adsorption has occurred, and the current hypothesis is that the SieA protein blocks entry of superinfecting phage DNA into the target cell. *Salmonella enterica* phage P22 SieA protein is the only system studied to date that is thought to block DNA injection by superinfecting short tailed phages, and in this report we examine phage P22 *sieA* mediated superinfection exclusion in more detail.

## RESULTS

### The phage P22 *sieA* gene is sufficient to block phage DNA injection

Prophages with a mutant *sieA* gene fail to exclude P22 (18, 24), and a high copy number plasmid carrying the P22 *sieA, orf59a* and *mnt* genes conveys strong resistance to P22 infection (25). In order to confirm that the *sieA* gene alone in low copy number is sufficient for superinfection exclusion we inserted it into the *S. enterica* LT2 strain UB-0002 chromosome by recombineering (see MATERIALS AND METHODS; bacterial and phage strains used are listed in Table 1). In the resulting strain (UB-2520), the inserted *sieA* gene replaces the *galK* gene and is in the same transcriptional orientation as the *gal* operon; the inserted DNA only includes the *sieA* gene and the putative P_sieA_ promoter (25) with 85 bps upstream of the *sieA* start codon and 22 bps downstream of the stop codon. Since there was no galactose in the growth medium, *sieA* expression was likely governed by P_sieA_. Figure S1 shows that the chromosomal *sieA* gene has little, if any, effect on cell growth at 37°C in rich medium. This *sieA^+^* strain was not lysed by P22 infection in liquid culture at a multiplicity of infection MOI) of seven, and the inserted *sieA* gene lowered the plaque forming ability of P22 by at least seven orders of magnitude (Figure 1). We note that loss of host *galK* function alone in the isogenic strain UB-2666 does not affect P22 infection. Thus, the *sieA* gene in one copy per *Salmonella* genome is sufficient to mediate robust superinfection exclusion.

**Figure 1.**
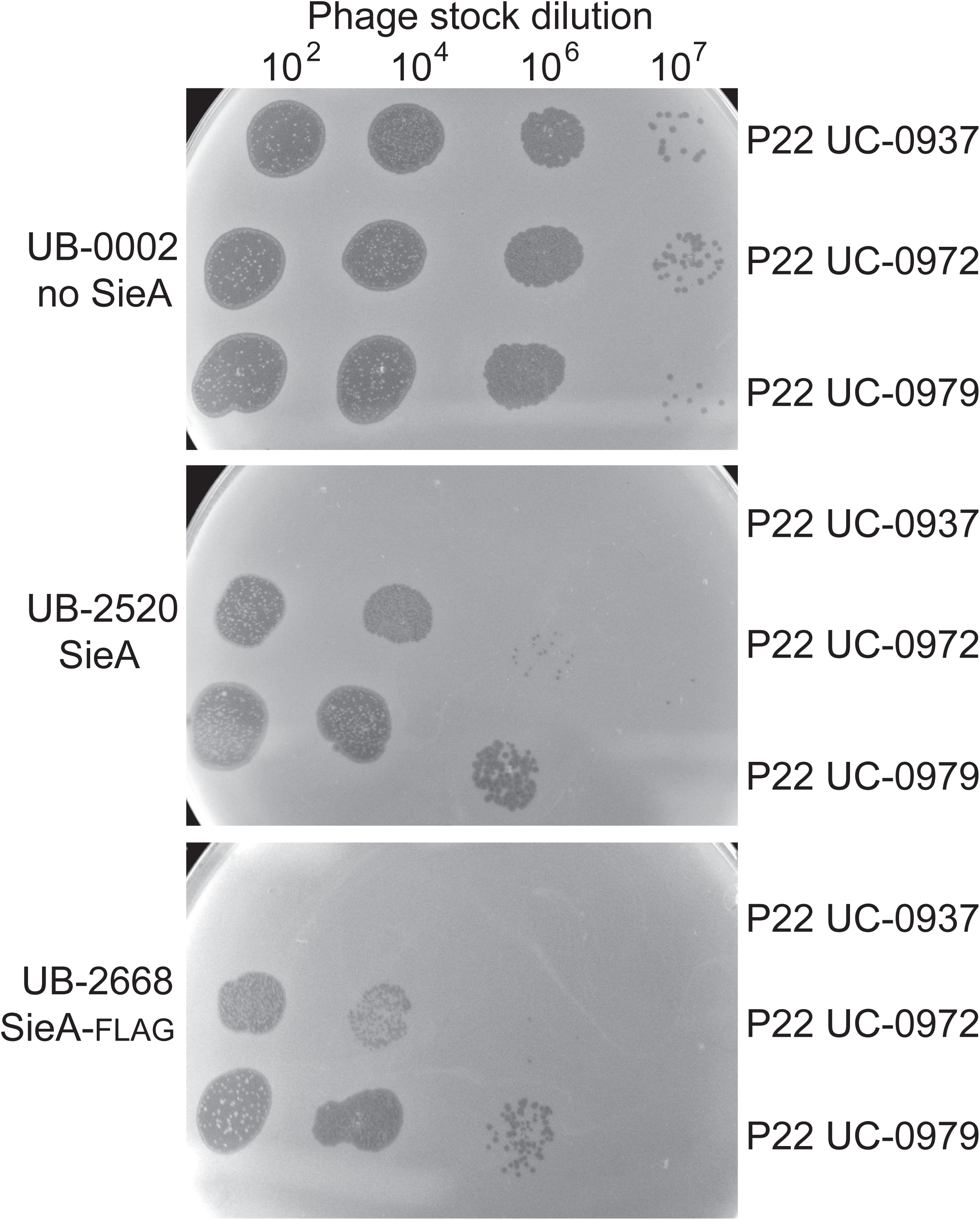
SieA superinfection exclusion. Soft agar containing the *sieA^+^* and *sieA^−^ S. enterica* lawn cells indicated at the right was allowed to solidify on L broth plates, and ten μL spots of parental “wildtype” P22 UC-0937, double mutant (P22 UC-0972) and triple (P22 UC-0979) mutant phages that overcome *sieA* exclusion (see text for mutant descriptions) were applied. The plates were incubated overnight at 37°C. Phage stocks were about 1010 PFU/mL, and they were diluted by the factors shown above. Ten microliters of undiluted P22 UC-0937 phage stock also showed no plaques on UB-2520.

**Table 1.**
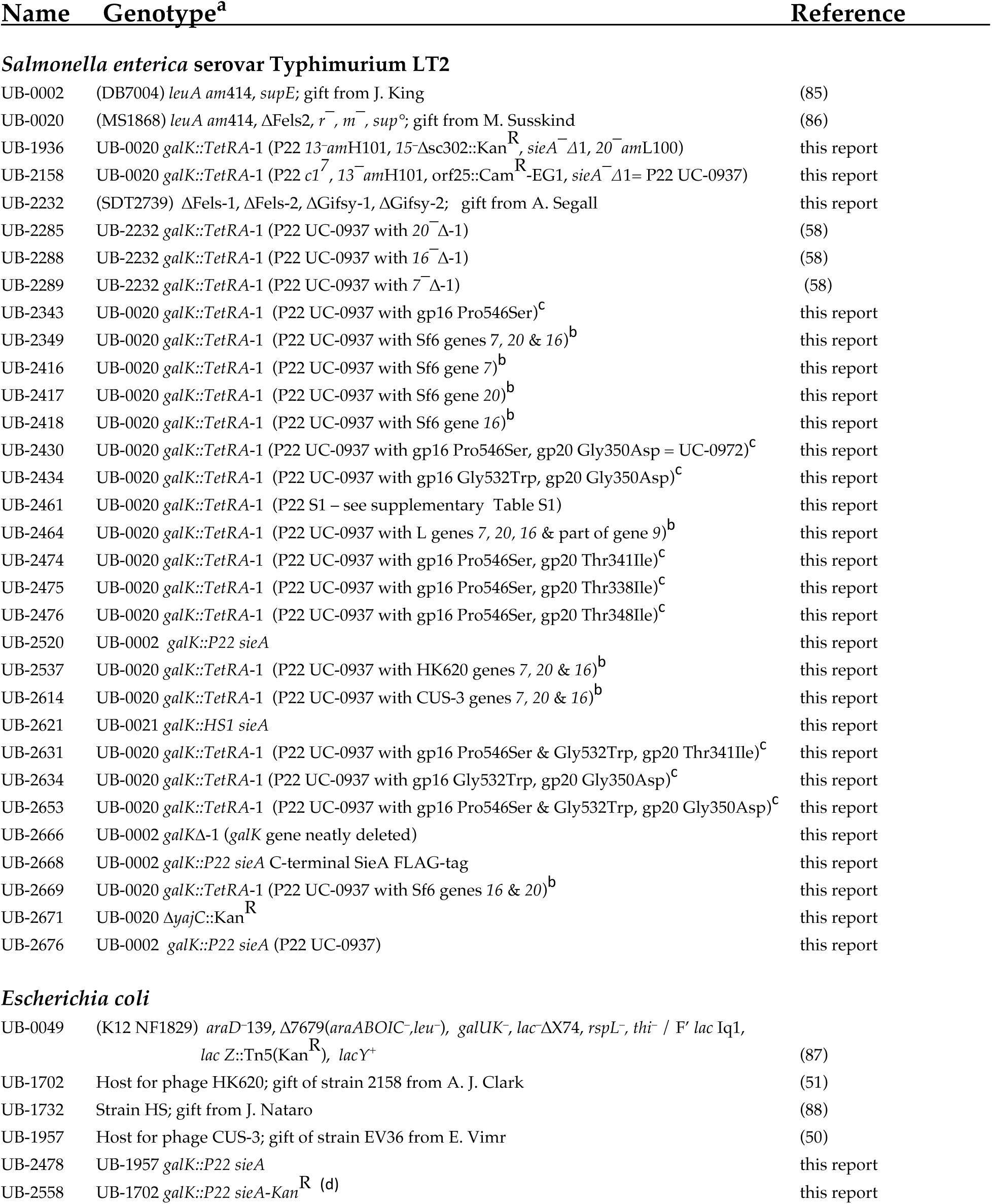
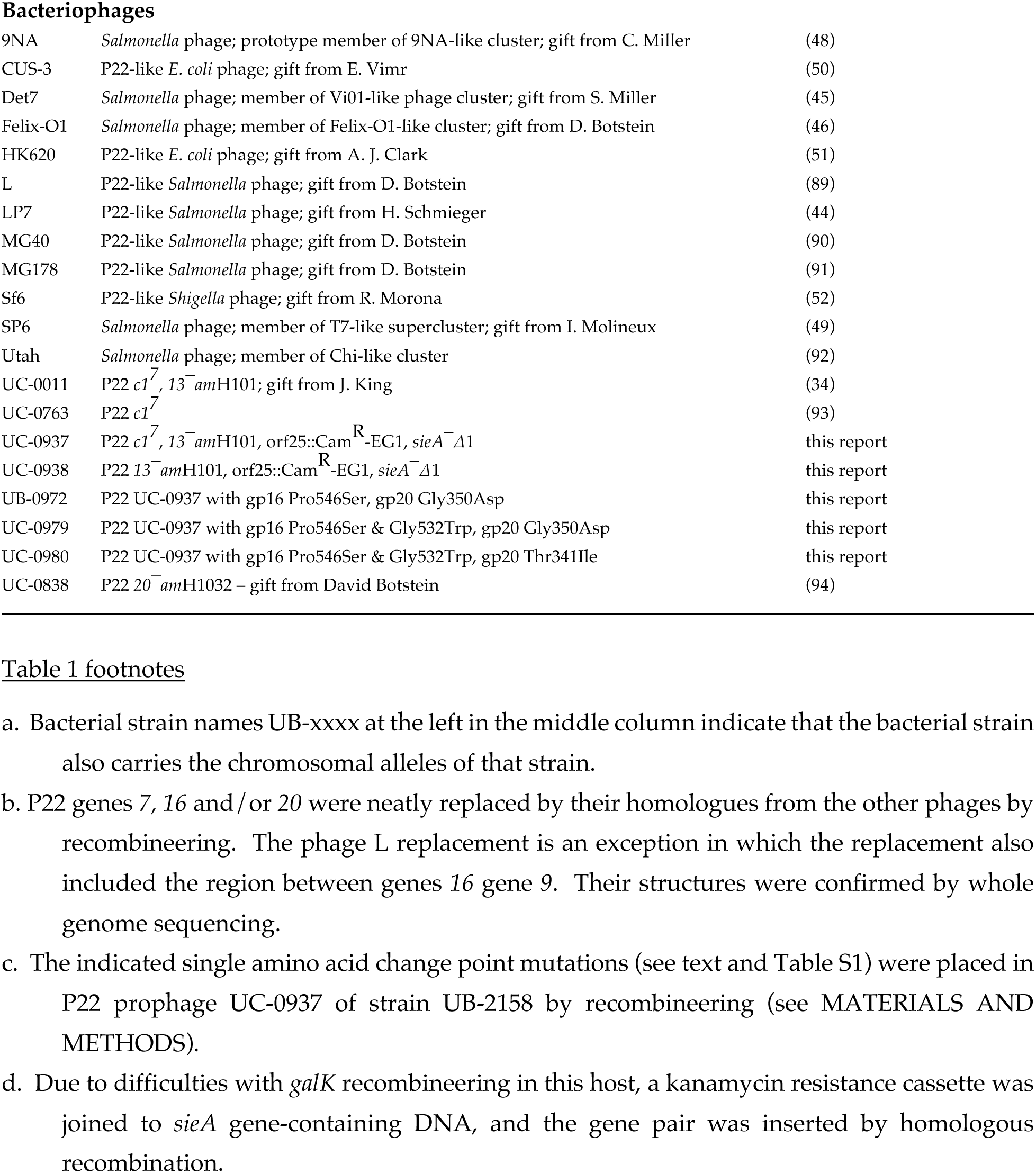
Bacteria and bacteriophage strains used in this study.

To confirm that under these conditions the superinfection block is very early in infection, we compared the frequency of lysogen formation by P22 lacking *sieA* in isogenic *sieA^+^* and *sieA^−^* host strains. Lysogenization should be a more accurate test for escape from SieA exclusion than is plaque formation since a rare successful single DNA entry event could give rise to a lysogen cell, whereas plaque formation requires successful progeny phage production in several successive rounds of infection. Isogenic cells with and without chromosomal *sieA^+^* genes (UB-2520 and UB-2666, respectively) were infected by P22 *c^+^*, *13^−^am*H101, *sieA^−^*, Cam^R^ (phage UC-0938; Cam^R^ is a chloramphenicol resistance cassette) at an MOI of 25 plaque forming units (PFU)/cell at 37°C in rich medium, and the resulting number of chloramphenicol resistant (lysogen) colonies was 10^5^-fold lower in the *sieA^+^* cells; the frequency of lysogens was about 25% for cells that do not carry the *sieA* gene. Furthermore, *sieA^−^* P22 (strain UC-0937; Table 1) prophages in isogenic hosts with and without the ectopic *sieA^+^* gene (UB-2676 and UB-2158, respectively) gave indistinguishable phage yields of about 10^10^ plaque-forming phage/mL after induction by addition of mitomycin C to 0.1 μg/mL. The differences between prophage induction and lytic growth are at the beginning of the growth cycle, thereby confirming that this is the stage at which SieA blocks infection under our conditions.

Finally, although the mechanism of K^+^ release upon infection is not fully understood, it has been shown to correlate very well with the successful delivery of DNA into the cytoplasm of target cells by a number of tailed phages including P22 (8, 26, 27). The current working model is that K^+^ escapes though the same channel through which the DNA enters the cell. Therefore, to test the ability of phage P22 to inject its DNA into *Salmonella* cells expressing SieA, a potassium selective electrode was used to measure K^+^ ion release into the surrounding medium from P22 infected cells. Figure 2A shows that P22 causes very little, if any, K^+^ release from the *sieA^+^* strain, while nearly all of the cellular K^+^ is released from the isogenic *sieA^−^* strain. We conclude that SieA blocks the injection of P22 DNA into *Salmonella* cells.

**Figure 2.**
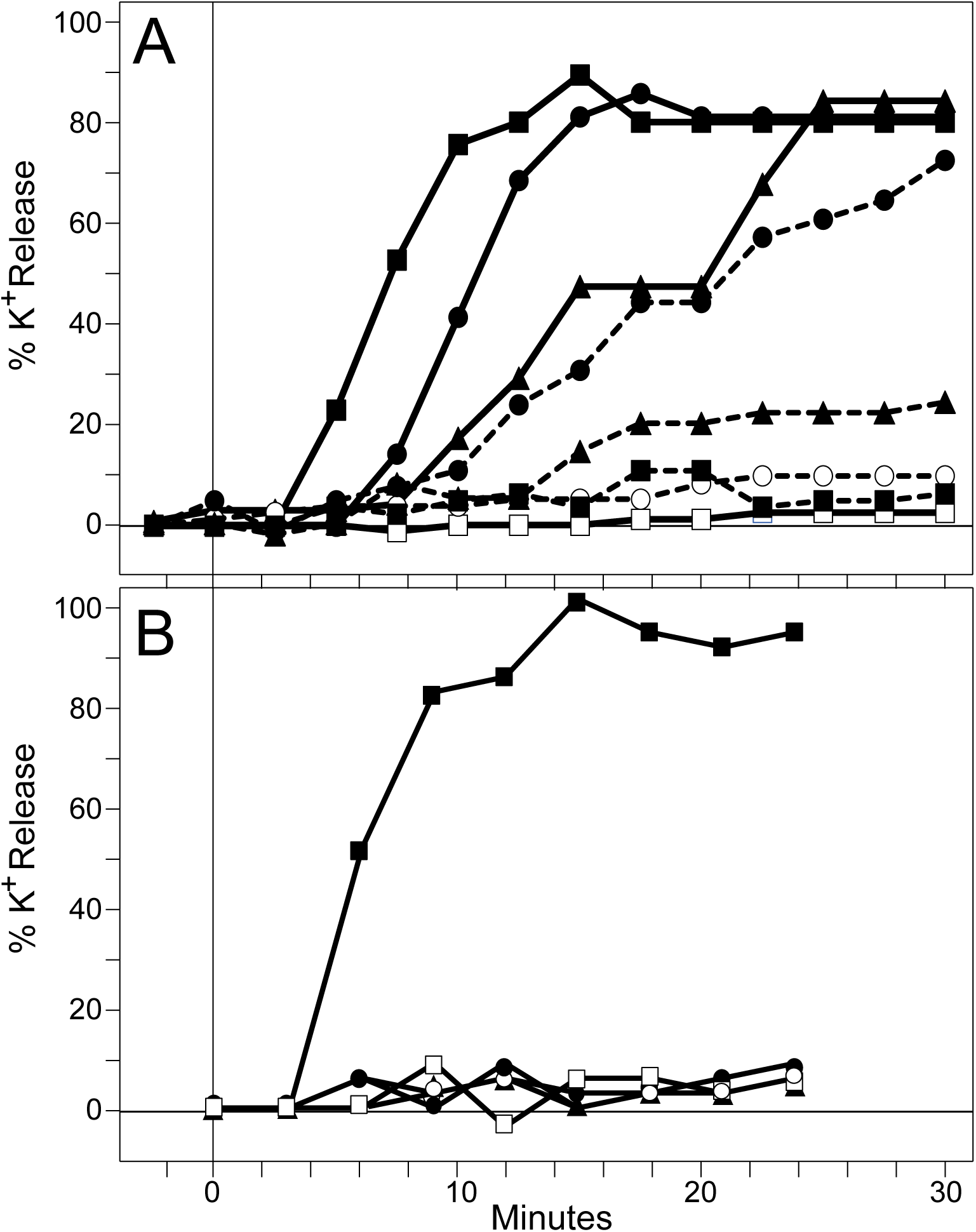
Potassium ion release after P22 infection. *S. enterica* strains growing in log phase were infected with P22 at 37°C, and K^+^ measurements (expressed as percent released relative to total K^+^ released after boiling for 10 min) were performed as described (27). Panel A. Potassium ion release by infectious P22 virions is blocked by P22 SieA protein. The *sieA^−^* UB-0002 host is represented by solid lines (–––) and the *sieA^+^* UB-2520 host by dashed lines (- - -). These strains were infected by parental P22 UC-0937 at MOI=5 (closed squares, -▪-), or with the SieA insensitive triple mutant P22 UC-0979 at MOI=5 (closed triangles, -l-) and MOI=10 (closed circles, -•-). Uninfected UB-0002 and UB-2520 are indicated by open squares (-□-) and open circles (-0-), respectively. Panel B. Lack of potassium ion release by P22 particles lacking E-proteins. The *sieA^−^* UB-0002 host was infected at MOI=5 by P22 UC-0937 (wildtype for genes *7*, *16* and *20*) is indicated by closed squares (-▪-), or by particles produced by induction of isogenic prophages deleted for one of the E-protein genes as follows: gene *7* deleted (UB-2289) (closed triangles, -l-); gene *16* deleted (UB-2288) (closed circles, -•-); or gene *20* deleted (UB-2285) (open squares, -□-). Uninfected UB-0002 is indicated by open circles (-0-). The particles in this panel were CsCl gradient purfied and relative concentrations of virion-like particles (which do not make plaques) were measured by quantitating the amount of coat protein in Coomassie Brilliant Blue stained SDS-PAGE gels, so that the same number of particles/cell was used in all cases. Full phage and host genotypes are given in Table 1.

### SieA presence and ejection protein (E-protein) absence have similar phenotypes

The phenotype of the P22 SieA block is similar to that of P22 virion-like particles produced by gene *7, 16* or *20* mutants. The protein products of these genes are released (ejected) from the virion during DNA delivery, and assemble into a transmembrane conduit that extends from the virion to the cytoplasm (28–31). Mutations that inactivate gene *7, 16* or *20* result in the formation of virions that lack the affected protein and that do not deliver their DNA into target cells properly (32–35).

Figure 2B shows that infection by E-protein deficient particles fail to release K^+^, which is similar to the infection of *sieA^+^* cells by wildtype P22 (above). Experiment using 7 or 30 particles/cell showed no measurable K^+^ release. Ejection of gp16 and gp20 are each independent of the presence of the other two E-proteins, and gp7 release is independent of the presence of gp20; however, gp7 is not released from particles that lack gp16 (33). Thus, release of gp20, gp7+gp16 or gp16+gp20 by *16^−^*, *20^−^*or *7^−^* particle infection, respectively, is not sufficient for release of K^+^ from the cell. In other words, all three E-proteins are required for K^+^ release and for DNA injection. The similar phenotypes that result from the SieA block or from the absence of any of the three E-proteins makes the idea that SieA may interfere with E-protein function during DNA injection an attractive hypothesis.

### SieA is an inner membrane protein

Although it has no obvious signal sequence, the 162 or 164 amino acid SieA protein (its start codon is uncertain) is predicted to be an integral membrane protein with three transmembrane domains (Figure 3A). Hofer *et al.* (25) reported that essentially all SieA protein is in the *Salmonella* cell membrane pellet fraction under presumed overproduction conditions, but they did not determine whether SieA is contained within the inner or outer membrane. In order to sensitively detect the SieA protein, we attached the FLAG epitope-coding sequence (36) to the 3’-terminus of the *sieA* gene to create strain UB-2668. This C-terminally tagged SieA has strong superinfection exclusion activity (Figure 1; Table 2). Inner and outer membrane fractions of isogenic strains UB-2520 (untagged, ectopically expressed SieA) and UB-2668 (FLAG-tagged SieA) were prepared according to method 1 of Thein *et al.* (37) (details in MATERIALS AND METHODS). Briefly, spheroplasts were prepared from cells grown to mid-log phase in L Broth at 37°C. After gentle lysis with Triton X-100, the outer membrane material was pelleted by centrifugation while the inner membrane material remained in the supernatant. The outer membrane fraction was further purified by centrifugation through a 35-40% OptiPrep density gradient. The proteins present in the whole cells as well as the inner and outer membrane fractions were separated by sodium dodecyl sulfate polyacrylamide gel electrophoresis (SDS-PAGE) and the FLAG-tagged band was identified by immunoblot analysis using monoclonal antibodies directed against the FLAG-tag (Figure 3B). In the immunoblot, these antibodies produced tagged SieA bands of approximately the same intensity in the whole cell extract and inner membrane fraction lanes of the FLAG-tagged strain UB-2668, but no such band was present in the outer membrane fraction. Parallel analysis of isogenic strain UB-2520 in which the SieA protein was not tagged showed no immunoblot band, confirming the specificity of the antibodies. Similar experiments with the somewhat less well documented membrane separation method of Sandrini *et al.* (38) also found that all SieA protein was in the inner membrane (data not shown). We conclude that SieA is an inner membrane protein.

**Figure 3.**
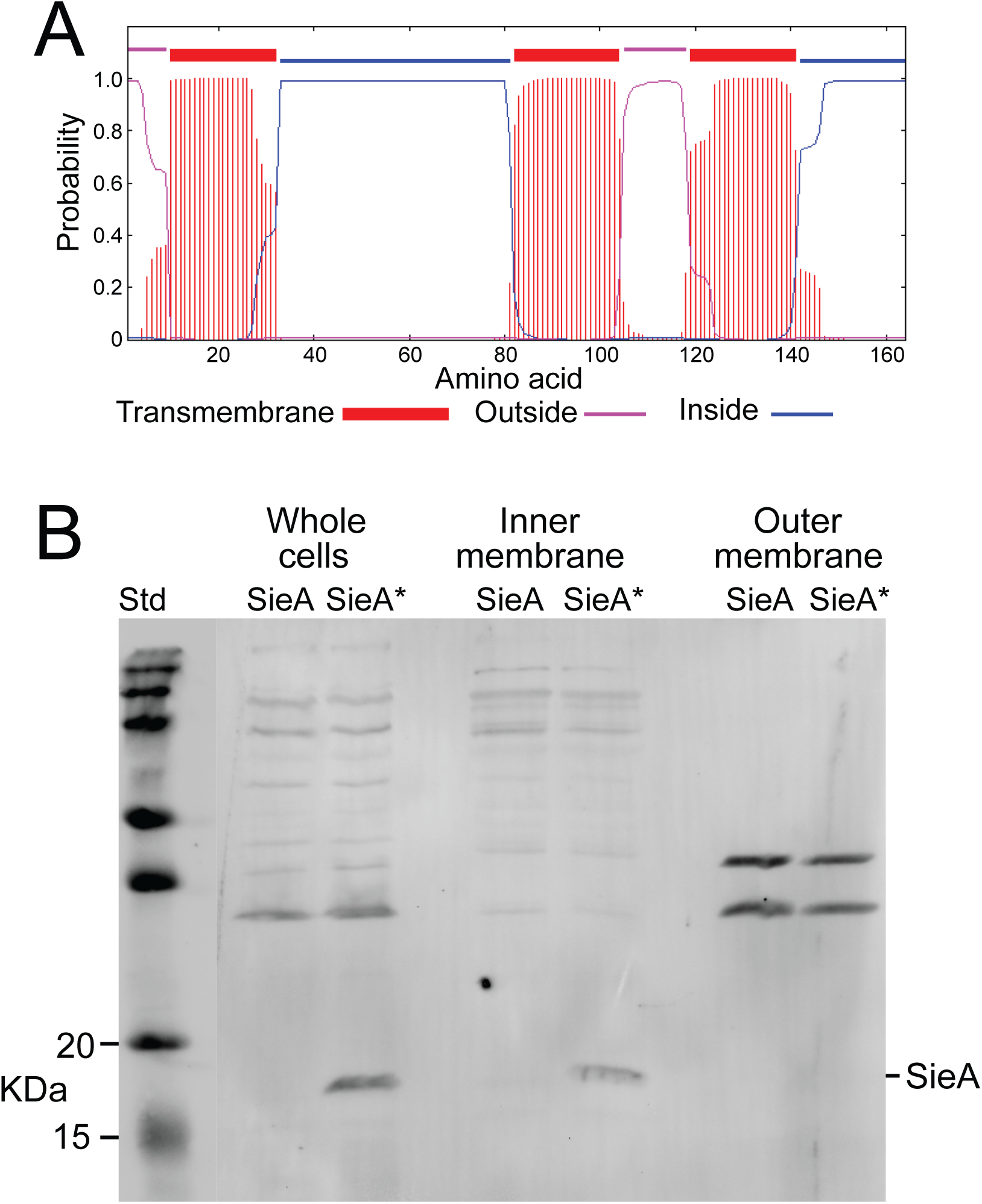
P22 SieA is an inner membrane protein. **A. P22 SieA protein membrane topology.** Membrane topology of SieA protein was predicted by TMHMM (54). **B. Cell fractionation.** A western SDS-PAGE immunoblot probed with an anti-FLAG monoclonal antibody conjugated to DyLight 800 4X PEG (ThermoFisher) is shown. The inner and outer membrane fractions of *S. enterica* strains carrying a P22 *sieA* gene (UB-2520) or a C-terminally FLAG-tagged P22 *sieA* gene (UB-2668) (marked with an asterisk, *) were isolated as described in MATERIALS AND METHODS. Samples were derived from equal numbers of cells of whole cell lysate, and inner membrane and outer membrane fraction proteins were separated by 15% SDS-PAGE. Precision Plus Protein Kaleidoscope (Bio-Rad) molecular weight markers are indicated on the left of each panel. Although some additional bands reacted with the anti-FLAG antibodies, the only difference between the tagged and untagged samples identifies the band that runs at the expected size of SieA protein (about 19 kDa) as the tagged-SieA protein band.

**Table 2.** SieA exclusion insensitive mutants of phage P22.

| <u>Lysogen</u><br><u>source<sup>a</sup></u> | <u>Phage genotype</u> | UB-2520<sup>b</sup><br>(P22 *sieA*<sup>+</sup>) | UB-0002<sup>b</sup><br>(P22 *sieA*-FLAG) | UB-2671<sup>b</sup><br>*yajC*<sup>-</sup> |
| --- | --- | --- | --- | --- |
| UB-2158 | P22 UC-0937 (parental “wildtype”) | <10<sup>-7</sup> | <10<sup>-7</sup> | <10<sup>-7</sup> |
| UB-2461 | Original escape mutant S1<sup>c</sup> | 0.04 | ND<sup>d</sup> | ND |
| UB-2343 | gene 16 Pro546Ser | <10<sup>-5</sup> | <10<sup>-7</sup> | ND |
| UB-2430 | gene 20 Gly350Asp, gene 16 Pro546Ser | 0.05 | 0.02 | ND |
| UB-2475 | gene 20 Thr338Ile, gene 16 Pro546Ser | 0.06 | 0.14 | ND |
| UB-2474 | gene 20 Thr341Ile, gene 16 Pro546Ser | 0.005 | 0.002 | ND |
| UB-2476 | gene 20 Ala348Thr, gene 16 Pro546Ser | 0.03 | <10<sup>-5</sup> | ND |
| UB-2634 | gene 20 Gly350Asp, gene 16 Gly532Trp | 0.1 | <10<sup>-5</sup> | ND |
| UB-2631 | gene 20 Thr341Ile, gene 16 Gly532Trp<br>and Pro546Ser | 0.5 | ND | <10<sup>-7</sup> |
| UB-2653 | gene 20 Gly350Asp, gene 16 Gly532Trp<br>and Pro546Ser | 0.7 | 0.3 | <10<sup>-7</sup> |

### P22 mutations that allow escape from SieA exclusion

Mutants of phage P22 that escape SieA-mediated superinfection exclusion have been reported but were not characterized (19). Our initial attempts at isolating such mutants by plating P22 *c1*^7^ phages (UC-0763), that were grown on *Salmonella* UB-0002, on a *sieA^+^* UB-2520 lawn generated rare plaque-forming phages at a frequency of less than one in 10^8^ infecting phages. Two of these were subjected to whole genome sequencing and were found to be hybrid phages that contain the P22 early region and the phage Fels-1 virion assembly genes. Fels-1 is a lambda-like prophage that is present in strains UB-0002 and UB-2520 (39, 40), and P22-Fels-1 hybrids have been observed previously (39, 41). Unlike those of P22, Fels-1 virions have lambda-like long non-contractile tails (41, 42). The fact that such hybrid phages successfully inject their DNA into cells expressing SieA protein lends yet more support to the notion that the P22 virion proteins are the target of SieA exclusion.

Initial attempts to isolate *sieA* escape mutants using P22 strains grown in a prophage-free host strain (UB-2232) did not give rise to any SieA resistant plaques. A mutagenized stock of clear plaque P22 *c1*^7^ (UC-0763) was therefore prepared by growing it on the prophage-free host in the presence of 5 μg/mL nitrosoguanidine. This concentration of mutagen gave the most *amber^+^* revertants of gene *13* in infections by P22 *c1*^7^, *13^−^am*H101 (UC-0011). When such mutagenized P22 *c1*^7^ phage stocks were plated on the *sieA^+^* UB-2520 host, approximately one plaque was obtained per 10^8^ applied phages. Eight plaques were picked from two independently mutagenized lysates and purified through three successive single plaque isolations on host strain UB-2232. These mutants made plaques with ≤6% efficiency on *sieA^+^* UB-2520 relative to isogenic *sieA^−^* UB-0002 (see for example original isolate S1 in Table 2). Their virions were purified and their DNAs were subjected to whole genome sequencing; they were found to carry between 7 and 25 new mutations (listed in supplementary Table S1), and none were sibling plaques as judged from their mutation content). Each of these eight phages carried more than one mutation in its E-protein genes *16* and *20*, and no other genes were universally affected in all mutants. All eight mutant isolates have a gp16 Pro546Ser substitution. Five of the eight also have a gp20 Gly350Asp substitution. The other three SieA escape phages carry one of following gp20 mutations instead of Gly350Asp: Thr338Ile, Thr341Ile or Ala348Thr (Tables 2 and S1). Their frequent presence in the escape mutants suggested that these E-protein changes were likely responsible for the observed partial SieA escape.

Recombineering methods were used to construct otherwise isogenic phages carrying these mutations (see MATERIAL AND METHODS). The prophage used for all recombineering in this study was P22 UC-0937. The construction of this phage is described in supplementary Figure S2: it carries a *13^−^amber* mutation that allows control of lysis after lytic growth; a *c1* mutation that lowers the frequency of lysogenization and so allows better lytic propagation without affecting the stability of the prophage state; a chloramphenicol resistance cassette that allows positive selection for lysogens; and a *sieA^−^* deletion that ensures robust synthesis of tailspike protein after prophage induction and obviates any potential complications due to SieA protein expression by the phage in the experiments presented below. None of these mutations affects prophage induction or progeny phage production. Perhaps somewhat surprisingly, the resulting gp16 Pro546Ser single mutant phage was unable to form any plaques on a *sieA^+^* host in spite of its presence in all eight escape mutants (it was not present in the starter phage in either of the two mutagenized stocks). Each of the four gene *20* mutations were also engineered into the prophage in combination with gene *16* Pro546Ser mutation. These double mutants form plaques on a *sieA^+^* host; however, like the original parental isolates, only ≤6% as efficiently as on a *sieA^−^* host (Table 2).

In order to isolate a P22 phage that forms plaques with a high efficiency on a *sieA^+^* host, the *16^−^* Pro546Ser, *20^−^* Gly350Asp double mutant (P22 UC-972) was propagated through multiple rounds of infection on a *sieA^+^* host as follows: A 25 mL L broth culture of *Salmonella* UB-2520 was grown to 5×10^7^ cells/mL at 37°C and infected with an isolated single plaque of P22 UC-972 from a *sieA^−^* (UB-0002) lawn, and the culture was shaken at 37°C for 5 hrs. Then 100 μl of this culture was diluted into a fresh 25 mL culture of *sieA^+^* UB-2520 and shaken again at 37°C for 5 hours. After nine such passages, a plaque on UB-2520 was identified whose phage plated with approximately the same efficiency on *sieA^−^* and *sieA^+^* hosts. This phage was subjected to whole genome sequencing, and it carried a single new mutation, a substitution of glycine to tryptophan at position 532 of gp16. To confirm this result, the Gly532Trp mutation was recombineered into the starting double mutant prophage to create a “clean” P22 *16^−^* Gly532Trp, *16^−^* Pro546Ser, *20^−^* Gly350Asp prophage (P22 UC-0979). This triple mutant phage makes plaques on a *sieA^+^* host with nearly wildtype efficiency (∼70%) as on a *sieA^−^* host, which is approximately 10-fold better than the starting double mutant (Table 2). Phage P22 *16^−^* Gly532Trp, *20^−^* Gly350Asp produced plaques at only about 10% efficiency, indicating that in this combination *16^−^* Pro546Ser still contributes to full escape function. A parallel triple mutant with a different gp20 mutation, P22 *16^−^* Gly532Trp, *16^−^* Pro546Ser, *20^−^* Thr341Ile (prophage UC-0980 in UB-2631) made plaques on a *sieA^+^* host with essentially the same full efficiency as P22 UC-0979. Since changes in E-proteins gp16 and gp20 allow escape from SieA exclusion, we conclude that SieA very likely blocks P22 DNA injection by interfering with the functions of these two proteins. The specific roles of gp16 and gp20 are discussed below.

Potassium ion release induced by infection with the *sieA* resistant triple mutant P22 UC-0979 is shown in Figure 2A. While the triple mutant released K^+^ to the same extent as the parental wildtype phage over the 30 min time course, the release of K^+^ was markedly slower for the mutant even in the absence of the host *sieA* gene. K^+^ was also released by the P22 UC-0979 triple mutant when a P22 *sieA* gene was present, but at MOI=5 it was slower and less extensive than release by the parental “wildtype” phage from a *sieA^−^* host. At MOI=10, the rate of K^+^ release by the triple mutant increased about 4-fold on the *sieA^+^* host (but not to the MOI=5 rate on the *sieA^−^* host) and allowed the extent of release to nearly reach the level of “wildtype” phage on *sieA^−^* host after 30 min (Figure 2A). Although the P22 UC-0979 triple mutant makes plaques with nearly wildtype efficiency, it appears to be partially impaired for K^+^ release and thus probably has partly impaired (slower?) DNA injection even on a *sieA^−^* host.

Bohm *et al*. (43) found the host inner membrane protein YajC to be essential for P22 DNA injection. It is thus a candidate for an inner membrane feature that the P22 injection apparatus (see below) contacts. However, neither of the P22 SieA resistant triple mutant phages (UC-0979 or UC-0980) made plaques on a *yajC^−^* host (Table 2) and, like wild type phages (see 43), their virions did not release potassium ions from *S. enterica* whose *yajC* gene had been deleted (UB-2671; data not shown). Thus, these *sieA* escape mutations do not bypass the YajC requirement.

### SieA target and host specificity

#### P22 SieA is specific for P22-like phages

Infection by P22-like short tailed *Salmonella* phages L, MG40 and MG178 is blocked by the *sieA^+^* gene expressed from a P22 prophage (19). We confirmed that these three phages as well as P22-like *Salmonella* phage LP7 (44) are unable to form plaques on *Salmonella* when an ectopically expressed *sieA* gene is present (strain UB-2520). On the other hand, we found that the following non-P22-like tailed phages make plaques with the same efficiency on *sieA^−^* (UB-0002) and *sieA^+^* (UB-2520) hosts: *Myoviridae* phages Det7 (45) and Felix-O1 (46), *Siphoviridae* phages ES18 (47) and 9NA (48), and *Podoviridae* T7-like phage SP6 (49). Susskind *et al.* (19) previously reported that Felix-O1 was not affected by SieA expressed from a prophage. Since SP6 is not affected, the P22 SieA protein does not exclude all short tailed phages and appears to be specific for P22-like phages.

#### P22 SieA is functional in E. coli

In order to test the ability of P22 SieA to exclude P22-like phages in another bacterial host species, *E. coli,* we inserted the P22 *sieA* gene into the chromosomes of the hosts of *E. coli* P22-like phages CUS-3 (type K1 strain EV36 (50)) and HK620 (type H strain 2158 (51)) resulting in strains UB-2558 and UB-2478, respectively. These two phages have tailspikes that bind different polysaccharides and therefore cannot infect each other’s host or *Salmonella*. CUS-3 and HK620 plate with efficiencies of ≤10^–8^ and about 0.3, respectively, on their *sieA^+^* host relative to the *sieA*^-^ host (Table 3). Thus, SieA from *Salmonella* phage P22 can function in *E. coli* since it excludes CUS-3 quite efficiently, but it did not function well against HK620 infection in its *E. coli* host. It was not clear from this experiment whether the failure to exclude HK620 was due to this phage’s insensitivity to P22 SieA exclusion or the failure of the *sieA* gene to be expressed or function in this host, but the hybrid phage experiments below strongly suggest that the former is true.

**Table 3.** P22 SieA protein functions in *E. coli*.

|  |  |  |
| --- | --- | --- |
| *S. enterica* host | UB-0002 <sup>a</sup><br>(*sieA*<sup>-</sup>) | UB-2520<br>(P22 *sieA*<sup>+</sup>) |
| P22 UC-0937 "WT" | 1.0 | <10<sup>-8</sup> |
| *E. coli* 2158 host | UB-1702<br>(*sieA*<sup>-</sup>) | UB-2478<br>(P22 *sieA*<sup>+</sup>) |
| HK620 WT | 1.0 | 0.33 |
| *E. coli* EV36 host | UB-1957<br>(*sieA*<sup>-</sup>) | UB-2558<br>(P22 *sieA*<sup>+</sup>) |
| CUS-3 WT | 1.0 | <10<sup>-8</sup> |

a. Values in the Table are plating efficiencies relative to the *sieA*<sup>-</sup> strain. Representative results are shown from three replicate experiments that gave very similar results.

#### Hybrid P22 phages that carry foreign E-protein genes

To further examine SieA target specificity and the idea that SieA inhibits E-protein function, we constructed and analyzed hybrid P22 phages whose E-protein genes were replaced by homologues from other P22-like phages. These genes have different names in each of the P22-like phages discussed here (e. g., 52); but, for simplicity, we refer to all their E-protein genes by the names of their P22 homologues. *Shigella* phage Sf6, *Salmonella* phage L, and *E. coli* phages CUS-3 and HK620 were chosen for these studies since their E-proteins are highly diverged (see below).

Recombineering methods were first used to construct a hybrid P22 prophage in which all three of its E-protein genes (*7, 16* and *20*) were replaced by the three Sf6 homologues (*Salmonella* strain UB-2349). This hybrid produced an essentially normal yield of functional phages upon induction with mitomycin C (Table 4 and Figure 4). However, when the individual P22 E-protein genes were replaced by their Sf6 homologues only the prophage with the Sf6 gene *7* homologue (UB-2416) gave a normal progeny yield, while induction of prophages with only Sf6 gene *16* (UB-2418) or gene *20* (UB-2417) did not produce plaque-forming phages. A prophage with Sf6 genes *16* and *20* (UB-2669) produced functional progeny (Table 4 and Figure 4). The nonfunctional virions produced after induction of P22 with only Sf6 gene *16* (UB-2418) contained Sf6 gp16 protein by SDS-PAGE analysis (data not shown); thus, their defect was not due to the inability of Sf6 gp16 to assemble into the particles, but rather due to its inability to function after assembly. In SDS-PAGE, Sf6 gp20 co-migrates with P22 coat protein and so could not be measured in particles that carry Sf6 gene *20*. The simplest explanation for the current observations is that gp16 and gp20 assemble into virions independently but must interact during DNA delivery, and P22 gp16 and gp20 cannot interact with Sf6 gp20 and gp16, respectively.

**Figure 4.**
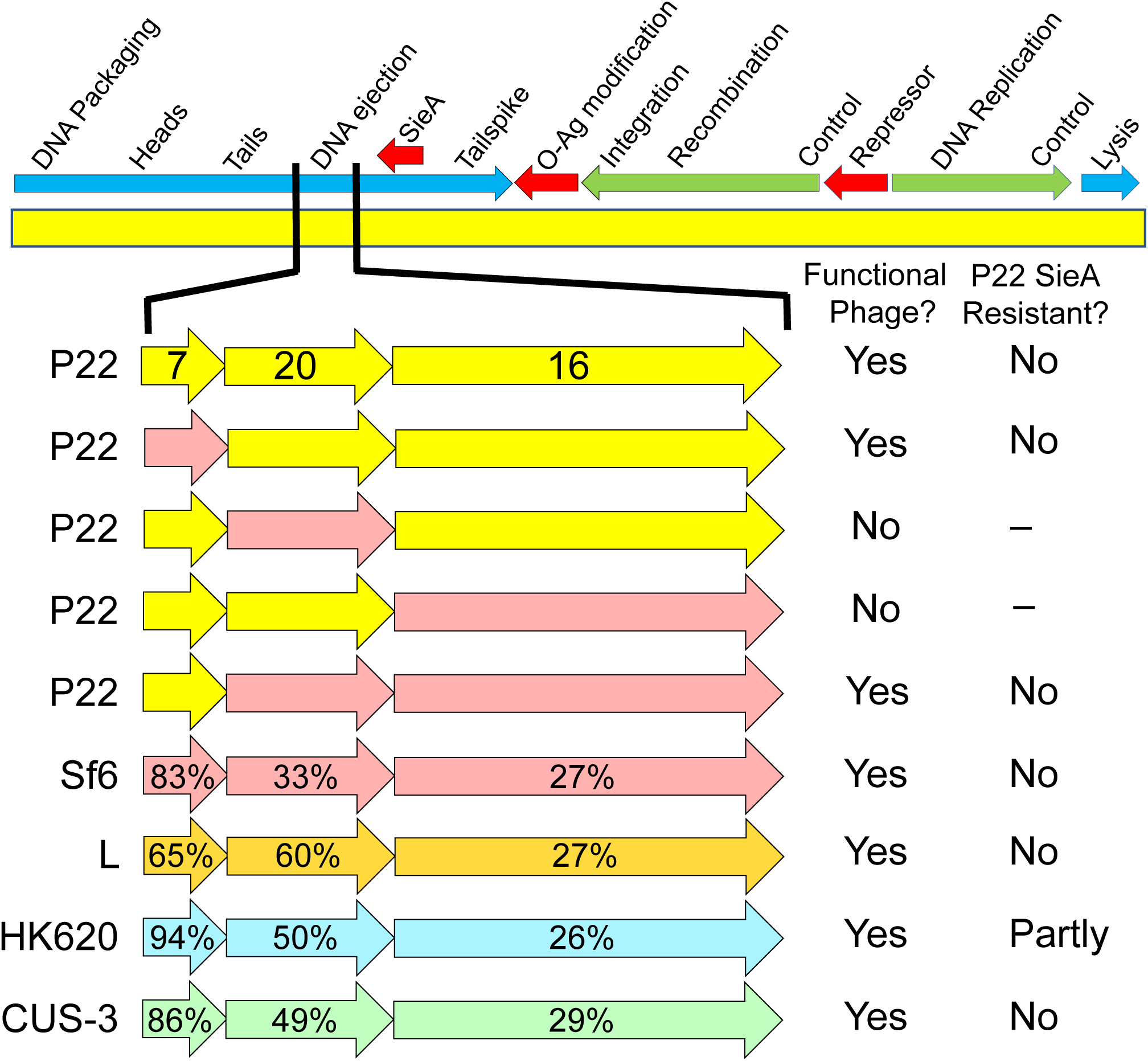
Phage P22 with foreign ejection protein genes. The phage P22 genome is diagrammed above with the location of the sieA genes indicated; red, green and blue arrows indicate regions expressed from the prophage and early and late during lytic growth, respectively. Below, replacement E-protein genes from phages Sf6 (strains UB-2349, -2416, -2417, -2418 and -2669), L (UB-2464), HK620 (UB-2537) or CUS-3 (UB-2614) are shown in different colors. P22 gene names are indicated on the P22 genes, and percent identities to their P22 homologues are indicated on the foreign genes. The viability and SieA resistance phenotypes are summarized on the right (see also Table 4).

**Table 4.** Functionality of P22 ejection protein hybrid phages.

| Ejection proteins<sup>a</sup> | Titer (PFU/mL)<sup>b</sup> |  |  |  | Phage source<sup>b</sup> |
| --- | --- | --- | --- | --- | --- |
|  | UB-0002<br>(*sieA*<sup>-</sup>) | UB-2520<br>(P22 *sieA*<sup>+</sup>) | UB-2668<br>(P22 *sieA*<sup>+</sup>FLAG) | UB-2621<br>(HS1 *sieA*<sup>+</sup>) |  |
| P22 gp7, gp20 & gp16 | 5x10<sup>10</sup> | <10<sup>4</sup> | <10<sup>4</sup> | <10<sup>4</sup> | UB-2158 |
| P22 gp20 & gp16, Sf6 gp7 | 1x10<sup>11</sup> | <10<sup>4</sup> | <10<sup>4</sup> | <10<sup>4</sup> | UB-2416 |
| P22 gp7 & gp20, Sf6 gp16<sup>c</sup> | <10<sup>4</sup> | ND<sup>d</sup> | ND | ND | UB-2418 |
| P22 gp7 & gp16, Sf6 gp20<sup>c</sup> | <10<sup>4</sup> | ND | ND | ND | UB-2417 |
| P22 gp7, Sf6 gp16 & gp20 | 5x10<sup>10</sup> | <10<sup>4</sup> | <10<sup>4</sup> | <10<sup>4</sup> | UB-2669 |
| Sf6 gp7, gp20 & gp16 | 2x10<sup>9</sup> | <10<sup>4</sup> | <10<sup>4</sup> | <10<sup>4</sup> | UB-2349 |
| L gp7, gp20 & gp16 | 1x10<sup>10</sup> | <10<sup>4</sup> | <10<sup>4</sup> | <10<sup>4</sup> | UB-2464 |
| CUS-3 gp7, gp20 & gp16 | 1x10<sup>11</sup> | <10<sup>4</sup> | <10<sup>4</sup> | <10<sup>4</sup> | UB-2614 |
| HK620 gp7, gp20 & gp16 | 2x10<sup>10</sup> | 3x10<sup>9</sup> | 4x10<sup>9</sup> | 2x10<sup>9</sup> e | UB-2537 |

P22 hybrid prophages were also constructed in which all three P22 E-protein genes were replaced by the phage L, HK620 or CUS-3 E-protein genes (strains UB-2464, UB-2537 and UB-2614, respectively). Upon induction, all three of these hybrids gave approximately normal yields of infectious virions (Table 4 and Figure 4), and SDS-PAGE analysis showed that each of these E-proteins was in fact incorporated into the hybrid virions (data not shown) as expected from their ability to form plaques.

Infection of *Salmonella* by the hybrid phages with all three Sf6, L or CUS-3 E-protein genes was efficiently blocked by P22 SieA; however, the parallel HK620 hybrid made plaques with about 10% efficiency when SieA was present compared to when it was absent (Table 4). The latter observation is consistent with HK620’s ability to form plaques with similarly reduced efficiency on its *E. coli* host when the *sieA* gene was present (above). The finding that P22 virions with HK620 E-proteins are only weakly sensitive to P22 SieA blockage strongly supports the idea that SieA protein interferes with E-protein function during DNA delivery into cells.

#### Diversity of *sieA* genes

Not all P22-like phages carry a *sieA* homologue. A search of the current public sequence database with BLASTp (53) found that 22 of the 66 P22-like phages whose genomes have been reported (as of June 7, 2023) carry *sieA* genes. These 22 phages all infect *Salmonella* and their SieA proteins are very close relatives of P22 SieA. Nonetheless, a more diverse set of *sieA*-like genes is present in bacterial genomes in the current public database. Most such *sieA*-like genes are present in *S. enterica* and *E. coli* genomes, but a few are in seven other bacterial genera – *Shigella, Serratia, Yersinia, Pectobacterium, Providencia, Proteus* and *Morganella* (all members of the family *Enterobacteriacae*). In all cases that were examined in detail, including over 20 each of the *Salmonella* and *Escherichia sieA*-like genes and 18 of those in the last seven genera, the *sieA* gene resides in a P22-like prophage and occupies a position similar to that of P22 *sieA*—between the homologues of P22 genes *16* and *9* (to be considered a *bona fide* prophage, the phage-like sequence must have been inserted into the host chromosome). There is substantial diversity among these prophage SieA proteins, and they are present as three main types—types A, B and C—that are only about 25% identical to one another in amino acid sequence (Figure 5). The *S. enterica* and *E. coli* prophage SieA proteins are either type A or B, and type C is currently limited to the *Providencia, Morganella* and *Proteus* genera.

**Figure 5.**
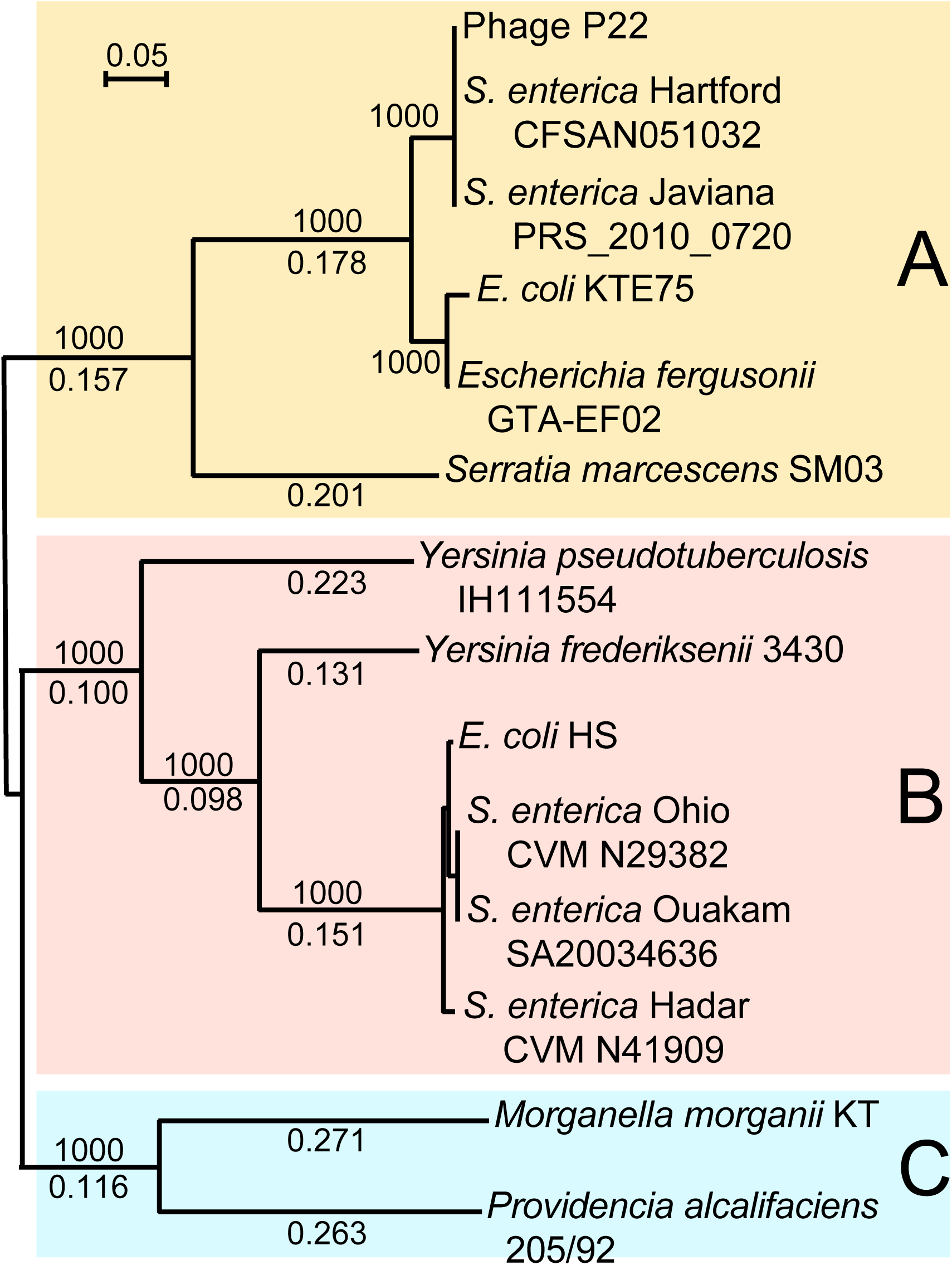
Neighbor-joining tree of representative SieA proteins. An unrooted Clustal neighbor-joining tree (67) of SieA proteins chosen to display the extent of their diversity is shown. The tree shows selected bootstrap values out of 1000 trials above the branches and fractional distances below the branches. Gray rectangles indicate the three sequence classes of SieA protein homologues. A bar representing a fractional distance of 0.05 is shown in the upper left. Supplementary Table S2 gives locus_tags for the SieA sequences.

The three SieA types have very similar transmembrane helix predictions by TMHMM analysis (54) (supplementary Figure S3). Types B and C have about 30 and 40 amino acid N-terminal extensions, respectively, relative to type A (P22 SieA is type A); and type C has about ten additional amino acids at the C-terminus. The extra N-terminal amino acids in types B and C contain an additional predicted transmembrane section and a putative N-terminal cytoplasmic region. Comparison of the three types shows weak sequence similarity scattered across the proteins with a somewhat more highly conserved motif, FnRYPnEAnnFNK, between P22 SieA residues 145-157 that is predicted to be on the cytoplasmic side of the membrane.

A type B SieA protein is encoded by the P22-like prophage HS1 in the *E. coli* strain HS genome (27, 55, 56). It is only 24% identical to P22 SieA protein. We examined the function of the HS1 *sieA* gene in more detail by PCR amplifying its DNA from strain HS (UB-1732; locus_tag EcHS_A0321; bp 325421-326206 in accession No. CP000802) and inserting it with 106 upstream bps into the same *galK* location in the *S. enterica* chromosome as the P22 *sieA* gene described above (strain UB-2621). *Salmonella* phage P22 (Table 4, top line) and P22-like phages L, MG40, MG178 and LP7 (not shown) were all fully excluded by HS1 SieA. In addition, similar to P22 SieA, HS1 SieA blocks infection of *S. enterica* by P22 hybrid phages with all three L, Sf6 or CUS-3 E-protein genes, but did not completely block the HK620 hybrid which again plated with about 10% efficiency (Table 4). In spite of their substantial amino acid sequence differences, the P22 and HS1 SieA proteins bestowed indistinguishable exclusion phenotypes in these experiments.

## DISCUSSION

### Inner membrane protein SieA blocks P22 DNA injection

We have shown that phage P22-encoded superinfection exclusion protein SieA is an inner membrane protein that specifically blocks injection of DNA into the cytoplasm of *S. enter*ica and *E. coli* cells from the virions of P22-like phages, and that no other phage-encoded proteins are required for this exclusion. Finally, we isolated P22 SieA escape mutants and hybrid phages in which changes in gp16 and gp20 overcome the SieA block. Three P22 E-proteins, gp7, gp16 and gp20, are present in P22 virions and are released from the virion after adsorption but before DNA injection (30, 33, 57, 58). Upon release from the virion, the E-proteins assemble into a hollow conduit for DNA passage from the virion through the host cell membranes, peptidoglycan and periplasm into the cytoplasm (28–30). This model for short tailed phage DNA injection is strongly supported by cryo-electron microscopic tomography of infecting short tailed phages T7 and P22 (31, 59–61). The observations here that (i) P22 infection by virions lacking E-proteins mimics the SieA block phenotype, (ii) P22 hybrids with phage HK620 E-proteins are partially resistant to SieA, and (iii) mutations that alter P22 gp16 and gp20 confer SieA resistance, suggest that SieA protein interferes with assembly of the E-protein conduit or with conduit function after its assembly.

*Lactococcus* phage Tuc2009 and *Streptococcus* phage TP_J34 also encode superinfection exclusion membrane proteins that block DNA injection into their Gram positive hosts (9, 10, 62, 63), and phage HK97 superinfection exclusion protein gp15 is also predicted to be an inner membrane protein in its Gram negative *E. coli* host (8). These three phages all have long-noncontractile tails, and their superinfection exclusion proteins appear to be specific for other long tailed phages. Our observation that P22 SieA is an inner membrane protein is the first for a short-tailed phage exclusion protein. Interestingly, it has been proposed that the long non-contractile tail tape measure proteins might form a transperiplasm DNA injection conduit for this type of phage (9, 64–66). The superinfection exclusion protein block can be overcome by alterations in the TP-J34 tape measure protein (9), and HK97 hybrids that have the phage HK022 tail tube and tape measure genes escape gp15 exclusion (8). Similarly, although it is very different from P22, the short tailed phage T7 also builds a conduit whose atomic structure has been determined (59–61). Its component proteins (T7 gp14, gp15 and gp16) have little recognizable sequence similarity to the P22 E-proteins. Thus, even though their component proteins have few obvious similarities in the above different phage types, the assembly of a transperiplasm DNA injection conduit seems likely to be a feature shared by all phages with short and long non-contractile tails.

### SieA and P22-like phage E-protein variation

We previously found that gene *16* and gene *20* are the among the most genetically variable parts of the P22-like virion assembly gene cluster; only the tailspike O-antigen receptor-binding domain is more diverse (55). It is perhaps surprising that very different E-proteins can successfully substitute for the P22 E-proteins in complete virion assembly and DNA injection (above). The E-proteins of the five phages examined here—P22, L, Sf6, HK620 and CUS-3—are quite divergent and highly mosaically related (55). The most highly conserved of these E-proteins are the gp7 homologues which range from 65% to 94% identical to one another in these five phages; it is not surprising that Sf6 gp7 can successfully substitute for the 82% identical P22 gp7 (Figure 5). The more diverse gp16 proteins are only 22-26% identical to one another, except for L and Sf6 gp16’s which are 75% identical. The gp20 proteins are 21-50% identical to one another, except for the L and P22 proteins which are 60% identical and the CUS-3 and HK620 proteins which are 98% identical. Since the CUS-3 and HK620 gp7 proteins and gp20 proteins are 90% and 98% identical, respectively, the ability of the HK620 E-proteins to mediate partial escape from SieA exclusion, while CUS-3 E-proteins do not, likely resides in their gp16 proteins which are only 26% identical. Among the gp16 and gp20 amino acids substituted in the SieA escape mutants (above), P22 gp16 Gly532 is conserved in all five of the above phages and P22 gp20 Thr338, Ile341 and Gly350 are conserved in three (by Clustal alignment (67)), suggesting that those SieA target residues may have conserved functions.

We showed that the type A SieA encoded by phage P22 and the type B SieA encoded by prophage HS1, which are only 24% identical, have indistinguishable effects on P22 and the four different P22 hybrid phages that have very different E-proteins. The interactions that mediate SieA exclusion must be quite robust since (i) three amino acid changes in the P22 E-proteins are required to fully overcome the plaque-forming block, and (ii) even with all three E-protein amino acid changes (above) the DNA injection rate is significantly slowed by the presence of SieA. In addition, the very different P22 and HS1 SieA proteins can both successfully exclude phages with distinct gp16 and gp20 proteins. If SieA interacts with the E-proteins as our results suggest, then it is difficult at present to understand how such interactions are maintained in the face of such extensive E-protein and SieA diversity. Understanding the details of these interactions will require further investigation.

### Role of ejection proteins in SieA-mediated exclusion

#### Gene *7* protein

There is no evidence that gp7 is directly involved in SieA exclusion. The relatively low extant diversity of gp7 suggests that it may interact with something more evolutionarily conserved than the host cell envelope, and an excellent candidate is the unusually highly conserved gp10 in the P22 virion’s tail (55). Indeed, gp7 may form the short tail extension from gp10 tail core seen in P22 virions that have spontaneously lost their DNA (68) and the phage virion proximal portion of the conduit (31).

#### Gene *20* protein

Purified phage Sf6 gp20 assembles *in vitro* into a 15 nm long tube that has an inner diameter of about 2.5 nm (69). Even though this may not be long enough to reach across the entire periplasmic space, which in *E. coli* is 18-25 nm (59, 70, 71), this structure probably forms at least part of the transperiplasm DNA delivery conduit, since the putative conduit is missing in P22 *20^−^* mutants (31). The gp20 tube is unlikely to be assembled with stacked globular gp20 units; its variable diameter fits much better with a model in which each gp20 polypeptide extends from one end of the tube to the other. This model also allows a facile explanation of how gp20 can tolerate the apparently random exchange of many *very different intragenic* mosaic modules (55) as follows: In a homo-multimeric structure built from parallel extended proteins that reach from one end of the structure to the other, most interactions will be short-range intra-subunit interactions between amino acids close to one another in the sequence and inter-subunit contacts between laterally aligned adjacent neighbors. Thus, any section of the protein could successfully be replaced by another sequence, even a very different sequence, as long as the new segment has the ability to maintain the side-by-side self-self interactions. If the gp20 tube is built from extended proteins, then the mutations in gp20 at codons 338, 341, 348 and 350 that help overcome the SieA block would be rather near an end of the gp20 tube (P22 gp20 is 471 amino acids long). We note that the C-terminal region of gp20 is essential for its function, since removal of the C-terminal 33 or 66 amino acids of P22 gp20 by nonsense mutations *am*H1032 and *am*L100, respectively, results in non-functional virions (sequences determined in this study).

#### Gene *16* protein

Thomas and Prevelige (72) reported that purified gp16 forms a rod-like assembly, but its structure has not been further investigated by high resolution methods. Wang *et al.* (31) reported that the transperiplasmic conduit is at least mostly formed in the absence gp16 and suggested that it resides in the inner membrane where the DNA would enter the cytoplasm. In agreement with this idea, Perez *et al*. (73) showed that gp16 associates *in vitro* with *Salmonella* membrane preparations and that gp16 facilitates *in vitro* DNA transport into liposome vesicles under certain conditions. In addition, a P22 mutation (tdx-1) that may affect circularization of generalized transducing DNA was reported to affect gp16 (74), which is consistent with an alteration in DNA entry. The gp16 SieA resistant mutations Gly532Trp and Pro546Ser probably do not affect the gp16 membrane association since the protein truncation experiments of Perez *et al.* (73) found that C-terminal amino acids 476-609 are not required for gp16 membrane association *in vitro*.

### Mechanism of SieA exclusion at the inner membrane

The presence of SieA protein in the inner membrane suggests that it interferes with DNA injection at that location. SieA could bind the conduit protein(s) to block conduit assembly or function. High MOI partially overcomes SieA exclusion of both infecting phage progeny and generalized transduction (18, 23, 24), suggesting that high levels of infecting E-proteins are able to saturate a constant and limited number of inhibitory host SieA protein molecules. This notion is consistent with transcriptomics data that show that *sieA* is not highly transcribed in the lysogen (75). However, *sieA* has a canonical ribosome binding sequence, and as such, with sufficient mRNA stability, could give rise high protein levels in the inner membrane. This in turn lends further credence to the idea that SieA interacts directly with P22 gp16 and gp20 transperiplasm conduit proteins.

An alternative hypothesis is that SieA occludes a required interaction between a host inner membrane component, likely YajC, and the distal end of the transperiplasm DNA delivery conduit. If YajC concentrations are in excess of SieA, which is consistent with transcriptomics data, then at high MOI there would be a higher probability that an infecting phage DNA conduit “finds” an unblocked YajC. Interestingly *sieA* is also upregulated during the course of prophage induction, suggesting that its exclusionary role becomes even more critical during phage particle production. It is possible that superinfecting P22 phages, whose genetic material and proteins would be generally indistinguishable from that of the prophage, could compete for cellular energy and protein synthesis machinery, thus thwarting the primary infection.

## MATERIALS AND METHODS

### Bacteria and phage strains

Bacteria and phage strains and their sources are listed in Table 1. *S. enterica* LT2 *supE* strain UB-0002 was used to propagate phage P22 strains, and *E. coli* UB-0049 was used to carry plasmids and as transformation recipient during plasmid construction. All plasmid and phage constructs were confirmed by determination of the sequence of the modified region after PCR amplification or by whole genome ILLUMINA sequencing at the University of Utah Sequencing Core Facility.

### Strain construction by recombineering

The *galK* recombineering methods used to modify phage P22 prophages were described in Padilla-Meier *et al.* (76) and Leavitt *et al.* (77). Briefly, in a *Salmonella* strain that carries the desired prophage (P22 UC-0937 here) and in which the native *galK* gene has been replaced by a TetRA tetracycline resistance cassette (78), the modification target was first replaced by a functional *galK* expression cassette (79) amplified by polymerase chain reaction (PCR) using primers that have ≥40 bp 3’-tail sequences with target homologies that program its precise insertion into the desired location by homologous recombination. A DNA of choice was then used to recombinationally replace the *galK* cassette by selecting for the concomitant loss of the *galK* gene by growth in the presence of 2-deoxygalactose (79). Plasmid pKD46 (80) was present to increase the efficiency of recombinational replacements, but was removed before any prophage induction experiments. Replacement DNAs for constructing P22 prophages with specific point mutations were created by using a synthetic double-stranded oligonucleotide containing the desired modification and ≥80 bp identity with the target. Replacement DNAs for constructing P22 prophages with E-protein genes from other phages were PCR amplified from DNA of the other phages with appropriate 3’-tails. The genome structures of these hybrid phages were confirmed by Illumina whole genome sequencing. In the Sf6 hybrid phages that did not give plaques (UB-2417 and -2418) the replacements, including both boundaries, were confirmed by PCR amplification and sequencing by Sanger (81) methods. In all cases except the phage L construct (UB-2464) the gene replacements were precise. In the latter case, the recombination that inserted the L E-protein genes included the first 23 codons of tailspike-encoding L gene *9* which is downstream of gene *16*, so this hybrid prophage contains the *7, 20* and *16* genes of phage L, the 836 bp between its gene *16* and gene *9* as well as this part of gene *9* (the first 23 amino acids of P22 and L tailspikes are identical, but there are four synonymous differences in this region).

To create ectopic *sieA* gene-carrying *S. enterica* and *E. coli* strains the bacteria were converted to *galK^−^* by recombineering replacement of the native *galK* gene by an appropriately PCR amplified P22 *sieA* gene (with or without C-terminal FLAG tag amino acid codons in the 3’-tail of the relevant primer) or HS1 *sieA* gene (82) followed by selection with 2-deoxyglactose.

### Nucleotide sequencing and sequence analysis

Sanger sequencing (81) of amplified PCR product DNAs and whole genome Illumina sequencing was performed by the High Throughput Genomics Core Facility, University of Utah. The latter utilized MiSeq 150-bp paired-end methodology with a 350-bp insert library. Quality controlled, trimmed reads were assembled into a single, circular phage sequence contig with >400-fold coverage using Geneious® 9.0.5 (83).

SieA protein homologue identification was carried out by examining the results of BLASTp (53) searches of the extant sequence database through the NCBI web site (https://blast.ncbi.nlm.nih.gov/Blast.cgi), and the neighbor joining tree was constructed with Clustal (67). Membrane topology of SieA protein was predicted by TMHMM (54) at the on-line site <http://www.cbs.dtu.dk/services/TMHMM/>.

### Cell fractionation, protein electrophoresis and immunoblot analysis

The inner membrane (IM) and outer membrane (OM) fractions of strains UB-2520 and UB-2668 were separated according to method 1 described by Thein *et al*. (37). The IM-containing supernatant was applied to a 4 mL Pierce detergent removal spin column (ThermoFisher Scientific) for removal of Triton X-100 and frozen at -20°C. The OM pellets were resuspended in a small volume of sterile distilled water by gentle shaking overnight at 4°C, applied to the top of 35-40% OptiPrep (Sigma) density gradients prepared in 30 mM MgCl_2_, 120 mM Tris-HCl, pH 8, and spun in a Sorvall MX120 ultracentrifuge at 4°C for one hour at 100,000 rpm. The gradients were fractionated and the resulting fractions were analyzed by SDS-PAGE after boiling for 3 minutes at 90°C in 3X sample buffer. The OM-containing fraction was stored at -20°C until Western blot analysis.

For Western blot analysis, equal amounts of the whole cell lysates as well as the IM- and OM-containing fractions of strains UB-2520 and UB-2668 were separated on a 15% acrylamide SDS-PAGE gel. Transfer to an Immobilon-PSQ membrane (Millipore) was carried out using a TE22 transphor electrophoresis unit (Hoefer Scientific) in 150 mM glycine, 20 mM Tris base, pH 8.3, 20% Methanol, 0.01% SDS overnight at 30 V at constant 0.1 Amp with stirring. The membrane was blocked with 10% nonfat milk in 20 mM Tris, pH 7.6, 150 mM NaCl for 1 hour with gentle shaking. The blot was washed for 5 minutes twice with TBS/T (20 mM Tris-HCl, 150 mM NaCl, pH 7.6 plus 0.01% Tween 20). The FLAG-tag monoclonal antibody conjugated to DyLight 800 4X PEG (ThermoFisher Scientific) was diluted to a final working concentration of 1:1000 in 1% nonfat milk in TBS/T. The membrane was incubated in the antibody solution for 2 hours at room temperature in the dark with gentle agitation and then washed for 5 minutes three times in TBS/T. The blot was rinsed and stored in TBS and visualized using a ChemiDoc MP imaging system (Bio-Rad).

### Analysis of transcriptomics data

Raw RNA-seq reads pertaining to wild-type P22 infection and prophage induction (75) were obtained from the NCBI SRA database (BioProject accession PRJNA737196). Raw reads were processed using BBDuk (v. 38.84) in Geneious Prime (2022.2.2) to remove adapter sequences and low quality reads. Individual processed read sets from either the infection or induction conditions were mapped to the *Salmonella enterica* subsp. *enterica* serovar Typhimurium str. LT2 (RefSeq accession NC_003197.2) and *Salmonella* phage P22 genomes (GenBank accession BK000583.1) using Geneious Mapper with default settings. RNA expression levels were obtained using the “Calculate Expression Values” function in Geneious with default settings, and differentially expressed genes were identified using the R package DEseq2 (84) within Geneious.

## ACKNOWLEDGEMENTS

We thank John Roth, Kelly Hughes, Donald Court, Anca Segall, Eric Vimr, Charles Miller, Miriam Susskind, Jonathan King, Stefan Miller, Host Schmieger, Ian Molineux, James Nataro, Renato Morona, A. John Clark and David Botstein for bacteria and phage strains. We also thank Lasha Gogokhia and Alexandra Heitkamp for help with strain construction, and Kristin Parent for helpful discussions.

---

**Table S1.** Changes in original mutants of phage P22 c1^7^ that overcome the SieA block.

| Mutant | <u>gp20</u> |  |  |  | <u>gp16</u> | Number of other changes |
| --- | --- | --- | --- | --- | --- | --- |
|  | Thr338Ile | Thr341Ile | Ala348Thr | Gly350Asp | Pro546Ser |  |
| S1 | – | – | – | + | + | 18 |
| S2 | – | + | – | – | + | 23, including 1 gene 20 synonymous |
| S5 | – | – | – | + | + | 5, including 1 gene 16 Gly191Ser |
| SA | – | – | – | + | + | 9, including 2 gene 16 synonymous |
| SB | – | – | – | + | + | 6 |
| SD | + | – | – | – | + | 9, including 1 gene 16 synonymous |
| SE | – | – | – | + | + | 11 including 1 gene 16 synonymous |
| SF | – | – | + | – | + | 14 including gene 7 Ala105Thr and<br>1 gene 16 synonymous |

**Table S2.** Prophage *sieA* gene locus_tags.

| <u>Bacterial Isolate</u> | <u>*sieA* homologue locus tag</u> |
| --- | --- |
| *E. coli* HS | EcHS_A0321 |
| *E. coli* KTE75 | A1UM_02908 |
| *Escherichia fergusonii* GTA-EF02 | VP22_0209700 |
| *Morganella morganii* KT | MU9_2893 |
| *Providencia alcalifaciens* 205/92 | HMPREF1563_1315 |
| *S. enterica* Hadar CVM N41909 | DE81_11120 |
| *S. enterica* Hartford CFSAN051032 | BUJ28_004415 |
| *S. enterica* Javiana PRS_2010_0720 | SEEJ0720_04091 |
| *S. enterica* Ohio CVM N29382 | DD58_22310 |
| *S. enterica* Ouakam SA20034636 | LFZ36_17085 |
| *Serratia marcescens* SM03 | A8A12_01925 |
| *Yersinia frederiksenii* 3430 | ERS008545_03773 |
| *Yersinia pseudotuberculosis* IH111554 | ERS008459_01168 |

**Figure S1.**
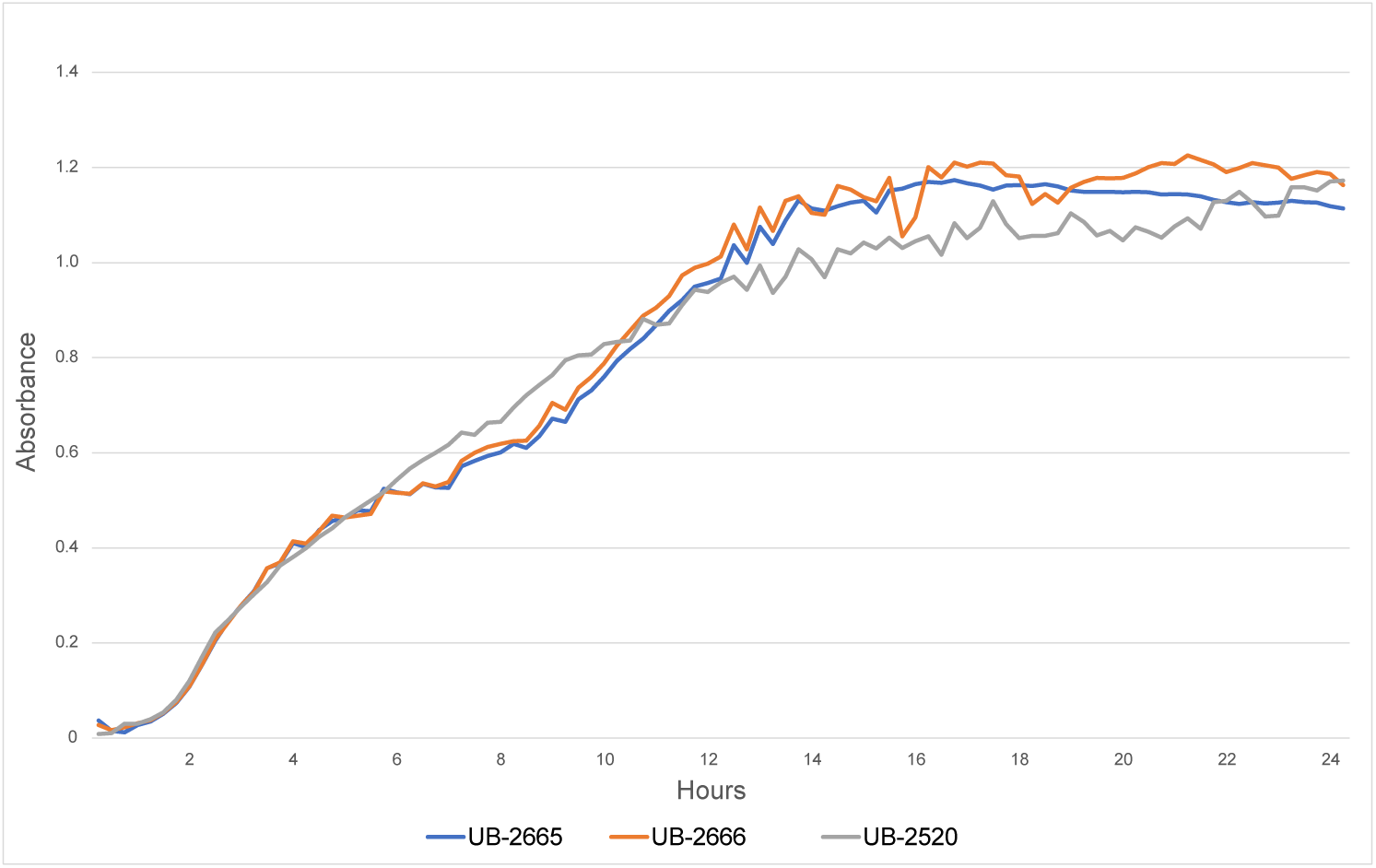
The P22 SieA protein has a minimal effect on *S. enterica* growth. Absorbance of bacterial cultures was followed with a Bioscreen C automated growth analysis system (Growth Curves USA, Piscataway, NJ) at 37°C in rich medium (L broth). *Salmonella* LT2 *galK^−^* strain UB-2666 and the isogenic *sieA^+^* UB-2520 have very similar growth curves. UB-2665, a restriction minus version of UB-2666 that is also missing the Fels2 prophage, shown for comparison, has a similar growth curve. Hofer *et al.* (1995) reported that expression of SieA from a *tac* promoter on a multicopy plasmid caused a growth rate reduction but did not quantify the effect. The Figure shows that the presence of the *sieA* gene in its chromosome has no strong effect on *S. enterica* cell growth at 37°C in rich medium. Thus, although it may be more detrimental in high levels, a single *sieA* gene has little effect on *Salmonella* growth under these conditions.

**Figure S2.**
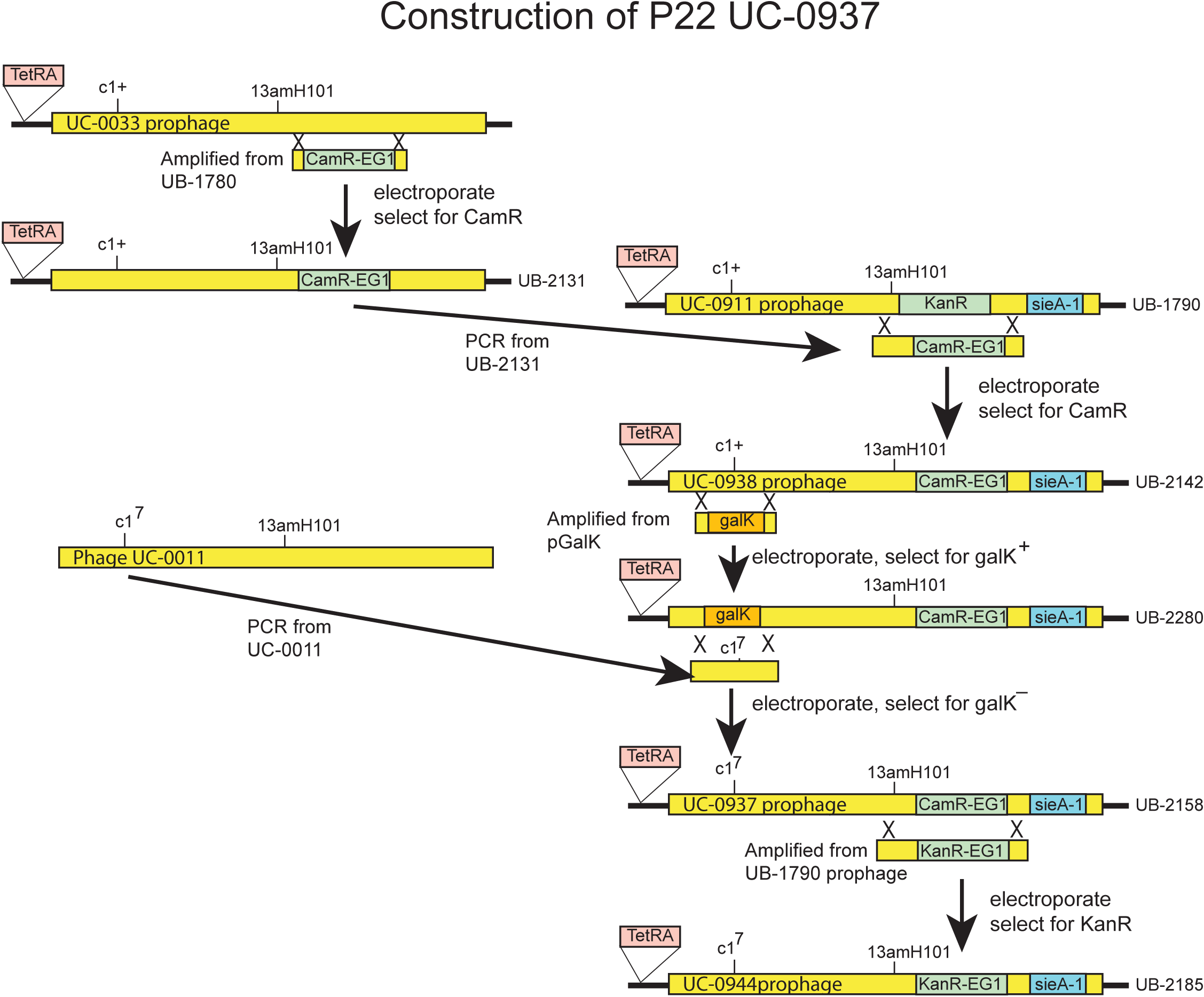
Construction of phage P22 phage UC-0937 - an improved P22 prophage recombineering substrate. We previously described hybrid phage constructs in parental prophage P22 *13^−^ am*H101, *15*^−^Δsc302::KanR, *sieA^−^Δ*1 (P22 strain UC-0911 prophage in bacterial strain UB-1790; Padilla-Meier *et al.*, 2012; Leavitt *et al.*, 2013a) in which the *13^−^amber* mutation allows control of lysis after lytic growth, the kanamycin resistance cassette allows positive selection for lysogens, and the *sieA^−^* deletion ensures robust synthesis of tailspike protein after prophage induction (see Adams *et al.*, 1985). In order to avoid the use of citrate plates for titering P22 phages (made necessary by the *15*^−^ deletion in phage UC-0911) and to allow more efficient lytic propagation of functional engineered phages, the genetic modification experiments here were performed with a modified prophage that also carries the *c1*^7^ clear plaque mutation and a chloramphenicol resistance cassette (orf25::CamR-EG1) that does not disrupt P22 gene *15* replaces the KanR cassette. The only P22 open reading frames replaced by the CamR cassette are orf25, orf80 and *pid*; the former two are not required for lysogeny or lytic growth but have not been studied further (Casjens *et al.*, 1989; Eppler *et al.*, 1991), and the latter is a nonessential gene that derepresses the host’s *dgo* operon in P22 pseudolysogen cells (Cenens *et al.*, 2013). The C1 protein stimulates establishment of the prophage state but is not required for its maintenance (Susskind and Botstein, 1978). This P22 *c1*^7^, *sieA^−^Δ*1, *13^−^am*H101, orf25::CamR-EG1 phage was named P22 UC-0937; it is present as the prophage in *Salmonella* strain UB-2158. Its construction is diagrammed in Figure S2, and the details of its construction and characterization are described in the following paragraphs. In Figure S2 genomes are not drawn to scale, and the pink TetRA insertion indicates the tetracycline resistance cassette that replaces (and thus inactivates) the host *Salmonella galK* gene. The chloramphenicol resistance (CamR) cassette of *Salmonella* strain UB-1760 was PCR amplified using primers with appropriate 3’-tail sequences, and this DNA was used to electroporate a P22 *13^−^am*H101 (phage UC-0033) lysogen, and chloramphenicol resistant cells were selected to give strain UB-2131. Homologous recombination of the 3’-tails of these primers with the phage P22 genome caused the amplified DNA to replace P22 bps 40205-40800 with the CamR resistance cassette (coordinates according to GenBank entry BK000583). The CamR cassette region was then amplified from the UB-2131 P22 prophage with several hundred bp of P22 sequence on both sides and the resulting DNA was used to recombinationally replace the kanamycin resistant cassette *15^−^*Δsc302::KanR in the P22 prophage of UB-1790 (Padilla-Meier *et al.*, 2012). The resulting chloramphenicol resistant, kanamycin sensitive strain was called UB-2142 (its plaque-forming prophage is P22 UC-0938). To place the *c1*^7^ clear plaque mutation (Levine and Curtiss, 1961) into the prophage of UB-2142, we first sequenced the *c1* gene from *c1*^7^ mutant P22 *c1*^7^, *13^−^am*H101 (phage UC-0011) and found that it carries one coding difference from the wild type gene, a G to T change at P22 bp position 32189 that alters the *c1* gene glycine GGG codon 52 to a tryptophan TGG codon. We then replaced the UB-2142 prophage’s *c1* gene with a *galK* gene as follows: The *galK* expression cassette was amplified with primers whose 3’-tails allow homologous replacement of P22 DNA between bp 31865 and 32974 (a region that includes the 3’-portion of the P22 *c1* gene, orf48 and the 5’-part of gene *18*, none of which are required for prophage maintenence). Electroporation with this DNA and selection for *gal^+^* cells resulted in a prophage in which *galK* replaces these P22 bps (strain UB-2280). About 1 kbp of DNA containing the *c1^7^* mutation (as well as orf48 and the missing part of gene *18*) was amplified from phage P22 UC-0011 (above) and used to recombinationally replace the *galK* gene in the prophage of UB-2280, resulting in *Salmonella* strain UB-2158 whose prophage was named P22 UC-0937. A kanamycin resistant version of phage UC-0937 was also made by recombinationally neatly replacing the UB-2158 CamR cassette with a KanR cassette amplified from UB-1790 DNA and selection for kanamycin resistance. This is strain UB-2185 and its prophage is phage P22 UC-0944. The P22 UC-0937 prophage in strain UB-2158 was characterized as follows: (i) Mitomycin C induction of UB-2158 growing in LB broth at a cell density of 2×10^8^/ml resulted in the release of about 10^11^ phage/ml (a phage yield similar to that of a *c^+^* P22 lysogen) that made clear plaques on UB-0002 but did not form plaques on UB-0001 (*i.e.*, it carries a clear mutation and an *amber* mutation in an essential gene). The induced lysogen lysed only after shaking with chloroform as expected of *13^−^* phage infections. (ii) Addition of tailspike protein did not increase the infectivity of phage produced after Mitomycin C induction of UB-2158, indicating that *sieA*^−^Δ1 allows good tailspike synthesis under these conditions and phage particles produced have the necessary complement of tailspikes. (iii) As expected, P22 UC-0937 is able to lysogenize *S. enterica*, but at a much lower frequency than *c^+^* phages; in an MOI=5 infection P22 UC-0937 formed chloramphenicol resistant lysogens on host UB-0002 about 10^–5^ as frequently as a parallel *c^+^* phage infection with P22 UC-0938. (iv) As expected, once formed, UC-0937 lysogens are stable and mitomycin C inducible (data not shown). (v) SDS polyacrylamide gel electrophoresis of purified P22 UC-937 virions showed the same protein bands as wild type P22 (data not shown). Finally, (vi) whole genome ILLUMINA sequencing of P22 UC-0937 showed that it contains the *sieA*^−^ϒ1 deletion (Padilla-Meier *et al.*, 2012), the *c1*^7^ mutation (above), the *13^−^am*H101 mutation (Rennell and Poteete, 1985), and the orf25::CamR-EG1 chloramphenicol resistance cassette. Not surprisingly, because of its complex history that involved crosses between various P22 strains that had long independent laboratory histories, some of which may not have been backcrossed after mutagenesis (Botstein *et al.*, 1972), the complete sequence of P22 UC-0937 showed that it also carries seven other point differences from the reported P22 sequence (Accession No. BK000583 bp coordinates) as follows: synonymous nucleotide changes G1658A in codon 559 of gene *1*, C35869T change in codon 14 of *ninF* and A39314G in codon 109 of gene *15*, an insertion of an A between bps C41200 and C41201 in the *orf80* gene fragment created by the chloramphenicol resistance cassette insertion, and nonsynonymous changes C20549T (Glu33Lys) in *gtrB*, C35023A (His92Asn) in *ninB*, G39142A (Arg52Gln) in gene *15*. Since P22-UC-0937 does not require citrate for efficient plaque formation, this mutant gene 15 is functional, and none of the other differences is expected to have an impact on lysogeny or lytic growth of the phage.

**Figure S3.**
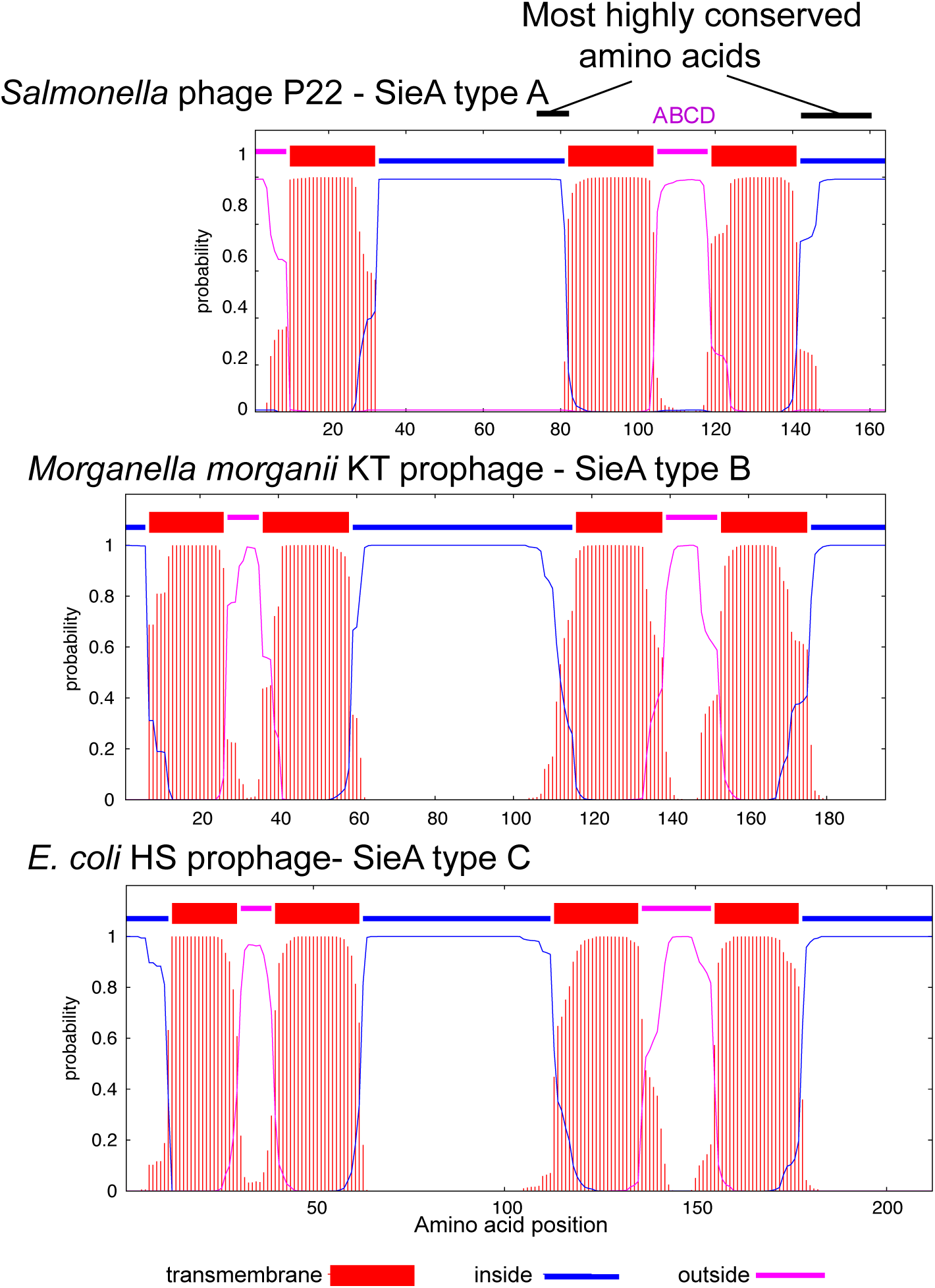
Predicted membrane topology of the three SieA sequence types. Membrane topologies of the three SieA protein types predicted by TMHMM (Krogh *et al.*, 2001). Note that types B and C have an N-terminal membrane spanning region that is not present in type A.

